# Genetic diversity and taxonomic issues in *Gastridium* P.Beauv (Poaceae) inferred from plastid and nuclear DNA sequence analysis

**DOI:** 10.1101/817965

**Authors:** Anna Scoppola, Simone Cardoni, Mariangela Pellegrino, Javier López-Tirado, Marco Cosimo Simeone

## Abstract

Incomplete taxonomic knowledge may seriously hamper biodiversity conservation efforts that are crucial in a context of global change. *Gastridium* P.Beauv. is a Mediterranean-Paleotropical member of the Poaceae family, inhabiting ephemeral grass habitats, whose species number and diversity are still imperfectly known. In order to progress towards a comprehensive taxonomic treatment of this genus, we examined patterns of DNA diversity in the four taxa (*Gastridium lainzii*, *G. phleoides*, *G. scabrum*, and *G. ventricosum*) that have been recently advanced by different authors, based on new morpho-ecological descriptors. We explored nucleotide sequence variation at two plastid (trnH-psbA, trnL-F) and one nuclear (ITS) DNA markers in 44 total individuals. Diversity data were treated with multiple statistical and phylogenetic tools, and integrated with available GenBank sequences of *Gastridium* and other closely related genera. Despite the limited variability detected, evidence of within-taxon genetic cohesion and estimates of molecular divergence comparable with those of species in the same subtribal lineage (Agrostidinae) were recovered. The identified plastid genealogies appeared congruent with a subdivision of the genus into (at least) three distinct entities, and coherent with collected morphological descriptors. Phylogenetic reconstructions with ITS were less corresponding to taxa identities, likely due to reticulation and polyploidization. Once placed in a broader taxonomic context, the investigated dataset produced plastid and nuclear tree topologies consistent with previous assessments, highlighting the overall little resolution of species and genera within Agrostidinae. In the plastid tree, a sister relationship between *Gastridium* and *Triplachne* was weakly supported. In the ITS tree, relationships among these genera were unresolved. The hypothesis of closely related but separately evolving lineages within *Gastridium* is discussed, suggesting a re-evaluation of its current assessment in taxonomic authorities to enhance our knowledge of the grass family, and assist future biodiversity surveys of key Mediterranean grassland ecosystems.

## Introduction

Grasslands (including rangelands, pastures, meadows, pseudo-steppes, and thermophile fringes) are important resources covering up to 48% of the whole Mediterranean region (Kyriazopoulos 2016). In their clearings and at their edges, they can incorporate little observed, neglected or transitional habitats rich in therophytes constituting an important asset of poorly known and endangered biodiversity (Blondel et al., 2014). Species in this biome respond to strict vegetation dynamics and are indicative of complex adaptive systems, including anthropogenic impacts (Clayton & Renvoize, 1986; Gallaher et al., 2019; Porqueddu et al., 2016).

*Gastridium* P.Beauv. is a Mediterranean-Paleotropical (Pignatti, 1982, Bolòs & Vigo, 2001) genus of the Poaceae family, represented by few, rather similar annual species inhabiting synanthropic, ephemeral habitats of Mediterranean shrubby pastures, garrigues, hedges, roadsides, and forest clearings. It is characterized by dense panicles with heteromorphic unifloral spikelets, each bearing unequal, slightly saccate glumes, laterally compressed and acute at the apex, with subterminal awn or unawned lemmas, and caryopses enclosed between the lemma and palea (Doğan, 1985; Scholz, 1986). Despite an overall morphological distinctness (Kellogg, 2015), resolving the taxonomy of *Gastridium* has been particularly challenging in the past, and its members have been alternatively included in other genera such as *Agrostis* L., *Milium* L., *Lachnagrostis* Trin., *Calamagrostis* Adans., and *Triplachne* Link (Scoppola & Cancellieri, 2019, and references therein). Today, it is placed within subfamily Pooideae Benth., tribe Poeae R.Br., subtribe Agrostidinae Fr. (Kellogg, 2015; Soreng et al., 2015a, 2017). Among the 11 genera of Agrostidinae (Soreng et al., 2017), the monotypic genus *Triplachne* [*T. nitens* (Guss.) Link; *Gastridium nitens* (Guss.) Coss. & Durieu] appears as the most closely related to *Gastridium* (Clayton & Renvoize, 1986; Quintanar et al., 2007; Saarela & Graham, 2010), although their precise affinities, and relation with the other genera are unclear (Saarela et al., 2017).

Species number and diversity within *Gastridium* is still an unsettled issue. According to Kellogg (2015) and Soreng et al. (2017), the genus comprises two species from Europe, North Africa and the Middle East: *G. ventricosum* (Gouan) Schinz & Thell., ≡ *Agrostis ventricosa* Gouan, and *G. phleoides* (Nees & Meyen) C.E. Hubb., ≡ *Lachnagrostis phleoides* Nees & Meyen. Both species are native to Italy; *G. ventricosum* has a wide mainland and insular distribution, and *G. phleoides* is currently known to occur on the Tyrrhenian coasts and Sicily (Bartolucci et al., 2018), although it is probably more widely distributed (Pignatti, 2017). Nevertheless, authors of Mediterranean flora consistently show evidence of additional morpho-ecological variation within the genus, suggesting the occurrence of further taxa, overlooked in major taxonomic treatments (e.g., Scholz, 1986; Valdés & Scholz, 2009). One of these entities is the Steno-Mediterranean *G. scabrum* C. Presl, recorded in Northwest Africa, East Spain, South France, Central and South Italy, Greece and the Aegean, Northwest Turkey, and West Asia (Bolòs & Vigo, 2001; Dimopoulos et al., 2013; Doğan, 1985, Feinbrun-Dothan, 1986; Pignatti, 2017; Pyke, 2008; Tison & De Foucault, 2014). It can be found on Mediterranean dry areas, pastures and abandoned fields, where the turf is broken by patches of crumbling soil (Scoppola & Cancellieri, 2019). A fourth taxon is *G. lainzii* (Romero García) Romero Zarco, ≡ *Gastridium phleoides* subsp. *lainzii* Romero García, a narrow endemic restricted to Southern Spain and Northern Morocco, growing in uncultivated vertisols (“bujeos”) and gravel road edges (López Tirado & Scoppola, 2017; Romero García, 2019; Romero Zarco, 2013). General poor knowledge of this complicated little genus results in units either synonymised (e.g., Romero García, 2019; Valdés & Scholz, 2009), or subordinated at subspecific ranks [e.g., *G. ventricosum* subsp. *scabrum* (C. Presl) O. Bolòs, *G. phleoides* subsp. *lainzii* Romero García], and in doubtful assignations of specimens in field inventories and herbaria (Scoppola & Cancellieri, 2019). The different treatments are also mirrored on the main sources of plant taxonomy information such as The Plant List (http://www.theplantlist.org/), Euro+Med (http://www.emplantbase.org/), and GrassBase (Clayton et al., 2006 onwards), whereas *G. lainzii* was only recently added to Tropicos (http://www.tropicos.org). Clearly, the taxonomy of *Gastridium* is still unresolved, and there is confusion over what names to apply and to how many species, preventing prompt and unambiguous detection of these taxa, with potential drawbacks on knowledge and conservation of the highly diverse and fragile communities where they occur (Lovatt, 1981; Pignatti, 2017). Frequent co-occurrence in the same plant community (e.g., *G. ventricosum/G. scabrum*, *G. ventricosum/G. phleoides, G. ventricosum/G. lainzii;* Lambinon & Deschâtres, 1994; López Tirado & Scoppola, 2017; Pyke, 2008; Romero García, 1996; Scholz, 1986) and the large, sometimes overlapping variation of some characters traditionally reported in literature (e.g., panicle shape, ligule length; Scholz, 1986; Jauzein, 2003) are the most likely obstacles to a resolved taxonomy. Recently, an enhanced combination of diagnostic characters has been provided, mostly based on some qualitative characters of florets, neglected in standard floras (Scoppola & Cancellieri, 2019), allowing a consistent circumscription of these taxa.

In modern taxonomy, the introduction of molecular methods and the use of DNA sequences have led to remarkable progress in the delimitation of species and their relationships (Hajibabaei et al., 2007; Rouhan & Gaudel, 2014), previously based only on shared morphological similarities (synapomorphies; Hennig 1965; Wiley et al., 1991). Molecular phylogeny is now considered a necessary complement to morphological studies (Simpson, 2010). In the light of this, the monophyly of Poaceae was first assessed with combined morphological and molecular analysis (Barker et al., 2001; Catalán et al., 1997), and later defined in an extensive phylogenetic framework (Bouchenak-Khelladi et al., 2008; Grass Phylogeny Working Group II, 2012; Saarela et al., 2018, Soreng et al., 2017). Nevertheless, despite many studies within Pooideae (e.g., Catalán et al., 2004; Hochbach et al., 2015; Persson & Rydin 2016; Quintanar et al., 2007; Schneider et al., 2012), diversity and evolutionary patterns in Agrostidinae remain poorly understood (Saarela et al., 2017). The wide application of plastid DNA sequences in species delimitation and phylogenetic analyses is due to some relevant operational features (availability of universal primers, large amounts of sequences in public databases usable for comparison), coupled with uniparental non-recombinant inheritance and mutation rates generally suitable for intrageneric inferences (Olmstead & Palmer, 1994; Saarela et al., 2017; Shaw et al., 2005; Soreng et al., 2015a). In turn, the higher mutation rate of the nuclear ribosomal internal transcribed spacer (ITS) region has made this marker the most commonly used for taxonomic and phylogenetic resolution below the genus level (Baldwin et al., 1995). However, several potential drawbacks can seriously affect the utility of the ITS spacer, causing experimental failures or incorrect reconstructions of species diversity and relationships (Álvarez & Wendel, 2003; Nieto Feliner & Rossellò, 2007). Among the many plastid loci available, the trnL-F region has been largely used, separately or in combination with ITS, in several major Pooideae lineages and genera (e.g., Amundsen & Warnke, 2012; Catalán et al., 2004; Chiapella, 2007; Hunter et al., 2004; Quintanar et al., 2007; Torrecilla et al., 2004; Torrecilla & Catalán, 2002). GenBank however, is also extremely rich in trnH-psbA sequences, a fast-evolving plastid region, of high relevance to resolve phylogenies and species identification in taxonomically complex plant groups (Kress et al., 2005; Pang et al., 2012).

Extensive phylogenetic/diversity studies at the intrageneric/intraspecific level have been performed on a few genera within the Agrostidinae subtribe (Amundsen & Warnke, 2012; Saarela et al., 2017). Other works on higher-level phylogenies (cf. Persson & Rydin, 2016; Quintanar et al., 2007; Saarela et al., 2017; Saarela et al., 2018) generally limited intrageneric sampling of *Gastridium* to one or two individuals of the major species (*G. ventricosum* and *G. phleoides*), preventing any inference on the genetic diversity of the genus and any attempt to identify evolutionary lineages. In this study, we performed a combined analysis of three markers from two independent genomic sources (nuclear: ITS; plastid: trnL-F, trnH-psbA) on an extensive *Gastridium* sample set to examine diversity patterns among the different taxa and clarify their systematic relationships. Our ultimate objective was to contribute to a better taxonomy of this genus, which is an essential tool for adequate knowledge and advanced conservation of biodiversity (Mace, 2004; Wheeler et al., 2004).

## Materials and Methods

### Plant material

Forty-four samples belonging to all Mediterranean *Gastridium* taxa were analysed (21, 15, 6 and 2 samples of *G. ventricosum*, *G. phleoides*, *G scabrum*, and *G. lainzii*, respectively). Of these, 30 individuals were freshly collected in wild populations, mostly located in peninsular Italy and Spain, and 14 samples were harvested from CLU, COFC, PORUN, UTV herbaria (acronyms according to Thiers 2018) and the U.S. National Plant Germplasm System (NPGS). Species identification was carried out according to Romero García (2019) and Scoppola and Cancellieri (2019). Vouchered herbarium specimens of collected plants were stored at the herbarium of the Tuscia University (UTV). Caryopses from NPGS and mature herbarium samples for which DNA extraction appeared unfeasible were germinated in the greenhouse. One-two fresh or dried accessions of grass species belonging to Agrostidinae [hereafter circumscribed according to Soreng et al. (2017); *Agrostis castellana* Boiss. & Reut.*, Polypogon monspeliensis* (L.) Desf., *Triplachne nitens* (Guss.) Link], or co-occurring in the same habitats [*Alopecurus aequalis* Sobol., *Briza minor* L., *Cynosurus echinatus* L., *Gaudinia fragilis* (L.) P. Beauv.] were also collected and included in the downstream analyses. Geographical location and additional information about the investigated samples are shown in Fig. 1 and Supplemental online Table S1.

**Fig. 1.**
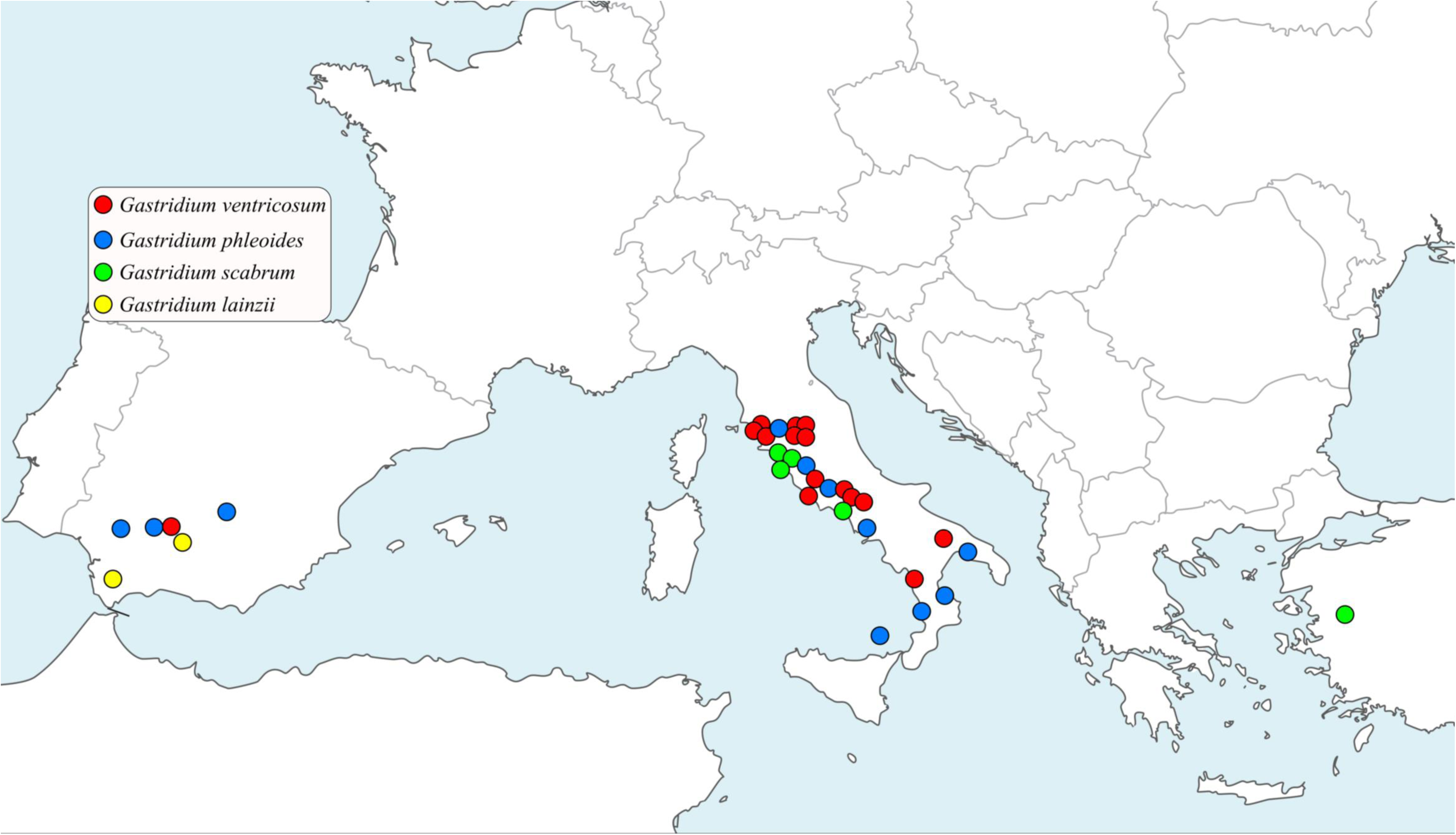
Geographical location of the investigated dataset

### Molecular analyses

Total DNA was extracted from silica gel-dried leaves and herbarium samples using NucleoSpin® Plant II (Macherey-Nagel) following the user manual. PCR reactions were performed with RTG PCR beads (GE Healthcare) in 25 μL final volume. The cpDNA regions (trnL-F, trnH-psbA) were amplified with the primers designed by Taberlet et al. (1991) and Shaw et al. (2007). The nuclear ribosomal internal transcribed spacer region (18S-ITS1-5.8S-ITS2-26S) was amplified with primers KRC (Torrecilla & Catalán, 2002) and ITS1 (White et al., 1990). Thermocycling conditions were 94 °C for 3 min, followed by 35 cycles of 94 °C for 3 sec, 53 °C (plastid regions) or 55 °C (nrITS) for 4 sec and 72 °C for 40 sec, with a final extension step of 10 min at 72 °C. PCR products were cleaned with GFX Illustra DNA Purification Kit (GE Healthcare); standardized aliquots were sent to Macrogen (http://www.macrogen.com) for sequencing, using the PCR primers of each marker. Electropherograms were edited with CHROMAS 2.6.5 (www.technelysium.com.au) and checked visually.

We also attempted a determination of the DNA content of three samples of each *Gastridium* taxon with flow cytometry. Given the lack of available specific standards and some problems encountered during the protocol set up, we consider our results only preliminary and present the adopted procedure and obtained data as a Supplemental online material S2.

### Phylogenetic analyses

MEGA 7.014 (Kumar et al., 2015) was used to build optimal multiple alignments and calculate the pairwise genetic distances for each (separate or concatenated) marker, and the mean estimates of inter- and intragroup molecular divergence. Diversity parameters for each marker were computed with DNASP 6.12.01 (Rozas et al., 2017). Median-joining (MJ) haplotype networks of every single and combined plastid region were inferred with Network 4.6.1.1 (http://www.fluxus-engineering.com/), treating gaps as 5^th^ state. The MJ algorithm was invoked with default parameters (equal weight of transversion/transition). The phylogenetic analyses of the ITS and concatenated plastid sequences were conducted with RAxML version 8.2.11 (Stamatakis, 2014) and focussed only on the sampled members of the Agrostidinae subtribe. The in-built GTRGAMMA model was selected with the „extended majority-rule consensus‟ criterion as bootstopping option (Pattengale et al., 2009), and 1000 replicates for assessing branch support (BS). A planar (Equal Angle, all standard parameters) phylogenetic network was produced with the Neighbor-Net algorithm implemented in SplitsTree4 (Bryant & Moulton, 2004, Huson & Bryant, 2006) to reveal complex phylogenetic signals of the ITS dataset; edge support was determined with non-parametric bootstrapping, and 1000 replicates.

Finally, we retrieved from GenBank (www.ncbi.nlm.nih.gov, accessed Aug. 2019) all available sequences of the most variable plastid marker (trnH-psbA) and the nuclear ITS of *Agrostis*, *Ammophila*, *Calamagrostis*, *Lachnagrostis*, *Podagrostis*, *Polypogon*, *Triplachne*, and *Gastridium*. These genera largely correspond to Agrostidinae p.p., i.e. one of the three major clades of this non-monophyletic subtribe, as currently circumscribed (Saarela et al., 2017). The extended datasets were analysed with MEGA and RAxML (see above) to compare estimates of molecular intra- and interspecific divergence and evaluate the phylogenetic relationships of the investigated dataset within the subtribe. *Briza minor* and *Briza humilis* were selected as outgroups in the phylogenetic analyses, based on Saarela et al. (2017, 2018) and Soreng et al. (2017). Vouchers of the *Gastridium* sequences retrieved from GenBank were requested to the hosting herbaria and examined to verify their taxonomic identity.

### Results

### Genetic diversity of *Gastridium*

Unambiguous electropherograms were obtained for 100% of samples with both plastid markers, whereas ITS sequence reactions failed in nine *Gastridium* and one *Gaudinia fragilis* samples, due to high density of double peaks and PCR contamination. Two *G. scabrum* individuals germinated in the greenhouse yielded pure plastid and nuclear DNA sequences, but the latter matched fungal ITS sequences on GenBank (100% identity) and were thus filtered. In total, 144 DNA sequences (121 belonging to genus *Gastridium*) were used for the downstream analyses. Table 1 shows the main diversity parameters exhibited by the individual markers and concatenated plastid regions in *Gastridium*.

**Table 1.**
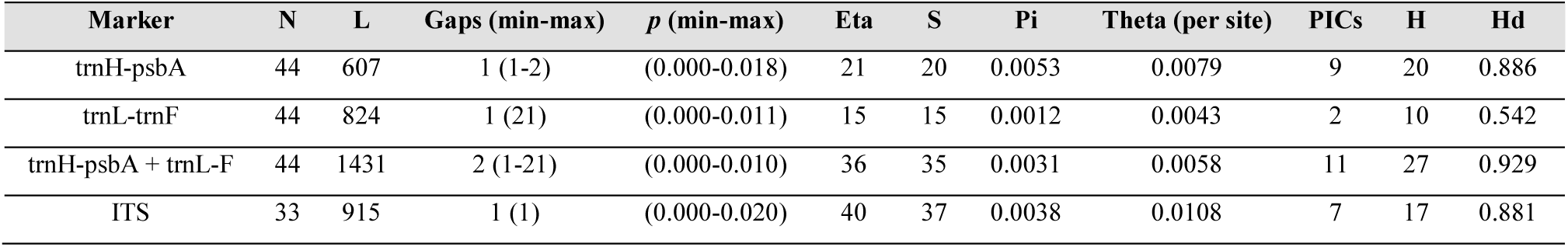
Markers‟ main diversity values in the investigated *Gastridium* dataset. N = number of samples; L = Aligned length (bp); Gaps = total number of recorded gaps (nucleotide range); *p* = uncorrected p-distance range; Eta = Total number of polymorphic sites; S = Number of polymorphic (segregating) sites; Pi = Nucleotide diversity; Theta = Nucleotide polymorphism (from Eta); PICs = Number of Parsimony Informative Characters; H = Number of haplotypes (with gaps included); Hd = haplotype diversity

TrnH-psbA was the most variable plastid marker, showing the highest number of polymorphic and segregating sites, the highest nucleotide diversity and estimated mutation rate, and one single gap, corresponding to a 1-2 nucleotide variation in a poly-A region. Overall variation resulted in a high number of parsimony informative characters and generated 20, highly diverse haplotypes. Nevertheless, the pairwise distance between sequences was moderately low. In contrast, the trnL-F region yielded lower diversity scores and generated half the haplotypes of trnH-psbA. However, its multiple alignment showed a 21-bp deletion that differentiated all samples of *G. scabrum* and *G. lainzii* from *G. ventricosum* and *G. phleoides*. This deletion was also detected in the other investigated genera, but not in *Gaudinia fragilis*. Longer insertions/deletions (range: 78-123 bp) were found in *G. fragilis*, *Cynosurus echinatus*, and *Alopecurus aequalis*. In total, the two concatenated plastid regions produced 27 highly diverse haplotypes. The nuclear ITS was the most variable marker, but only a small proportion of the variation encountered was scored as parsimony informative. The generated plastid haplotype and nuclear ITS variant lists (Supplemental online Table S3) showed only two trnL-F haplotypes shared between species pairs (*G. ventricosum/G. phleoides*, and *G. scabrum/G. lainzii*), whereas no trnH-psbA, concatenated haplotypes or ITS variants grouped members of different species (or genera). For the most part, the plastid haplotypes and ITS variants were unique or common to 1-3 individuals; only one plastid haplotype and one ITS variant were shared by more than ten individuals, all belonging to *G. ventricosum*.

The mean divergence calculated within and among the four *Gastridium* taxa (hereafter defined intra- and intergroup, respectively) with the two most variable (trnH-psbA and ITS) and the two concatenated plastid markers (trnH-psbA + trnL-F) are presented as Supplemental online Table S4. With all markers, *G. scabrum* exhibited mean intragroup divergence values (0.001, 0.003 for the plastid and nuclear regions, respectively) always lower than every mean intergroup estimate (0.002 - 0.007, and 0.005 - 0.009, respectively). In contrast, the intragroup mean divergences of *G. lainzii* were identical (0.003 at the ITS marker, 0.002 at the combined plastid regions) or lower than the intergroup values scored with *G. ventricosum* and *G. scabrum*. *Gastridium phleoides* was the most diverse and divergent, and scored a lower intergroup mean divergence of the ITS sequences with *G. ventricosum* and *G. lainzii* (0.003 - 0.005 vs. 0.008). Similar (intra-) and lower (interspecific) mean ranges of divergence were observed in other Agrostidinae genera with large occurrence of trnH-psbA and ITS sequences downloaded from GenBank (e.g., *Agrostis*, *Calamagrostis*, *Polypogon*; Supplemental online Table S4).

### Phylogenetic reconstructions

The single trnL-F and trnH-psbA haplotype networks (Supplemental online Figure S5a, b) provided little resolution of the *Gastridium* dataset. Two main groups, separated by two mutations and one median vector could be identified with trnL-F. The first one included haplotypes of *G. ventricosum* + *G. phleoides* (differing by 1-2 mutations, with the only exception of an isolated *G. phleoides* haplotype), and the other included *G. scabrum* + *G. lainzii*. The trnH-psbA network separated all the haplotypes produced in the four *Gastridium* taxa but was less resolved. The haplotypes of *G. phleoides* were characterized by a large diversity, with up to five mutations and a wide occurrence of median vectors separating most haplotypes. In both networks, haplotypes of *Triplachne nitens*, *Polypogon monspeliensis*, and *Agrostis castellana* were the least divergent from the *Gastridium* dataset; all other genera differed by >20 mutations.

The concatenated plastid haplotype network showed a better resolution (Fig. 2). Sequences were grouped in three major lineages separated by at least 2-3 mutations and 1-2 median vectors, accommodating (1) seven haplotypes produced in *G. ventricosum*, all differing by 1-2 mutations, (2) *G. scabrum* (four haplotypes) + *G. lainzii* (two haplotypes) differing by a single mutation, and (3) *G. phleoides* with fourteen haplotypes differing by 1-6 mutations. Eight mutations and two median vectors separated the *Gastridium* dataset from the haplotype of the closest genus (*Triplachne*).

**Fig. 2.**
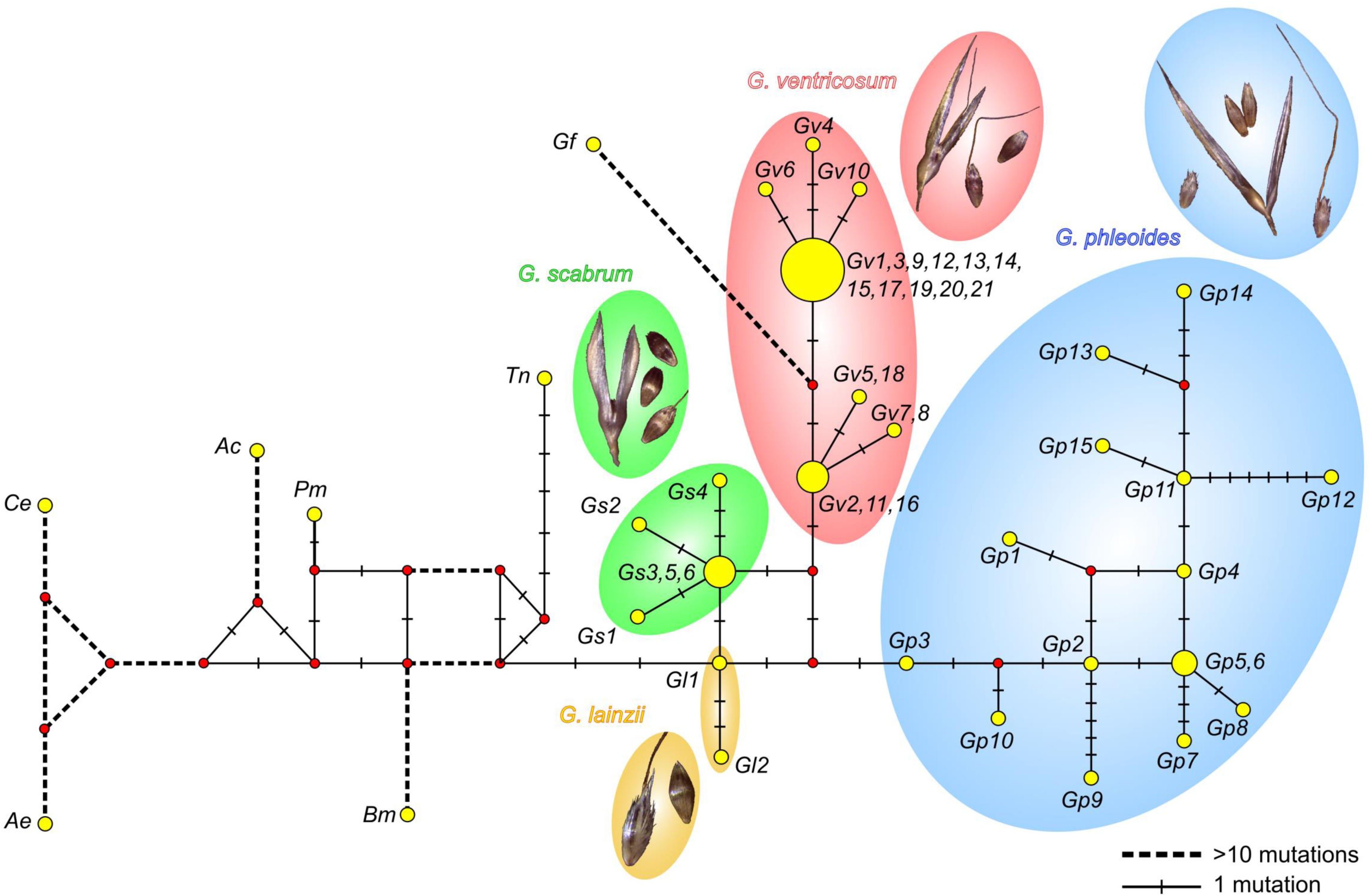
Median Joining haplotype network of the trnH-psbA + trnL-F concatenated regions. Coloured ellipses circumscribe *Gastridium* taxa (colours as in Fig.1). Different line thickness indicates the relative number of mutations. *Ac: Agrostis castellana, Ae: Alopecurus aequalis, Bm: Briza minor, Ce: Cynosurus echinatus, Gf: Gaudinia fragilis, Pm: Polypogon monspeliensis, Tn: Triplachne nitens, Gv: Gastridium ventricosum, Gp: G. phleoides, Gs: G. scabrum, Gl: G. lainzii*

Figure 3 shows the RAxML tree of the concatenated cpDNA regions. Agrostidinae samples split in two well-supported clades, with *Agrostis castellana* + *Polypogon monspeliensis* sister to a second clade including *Triplachne nitens* and all *Gastridium* samples. All *Gastridium* sequences were arranged in a moderately supported clade (BS = 78). Although the obtained sample grouping appeared largely coherent with taxa identities based on the used morphological descriptors (but less with the ecological patterns of distribution), resolution among the four taxa was unsupported.

**Fig. 3.**
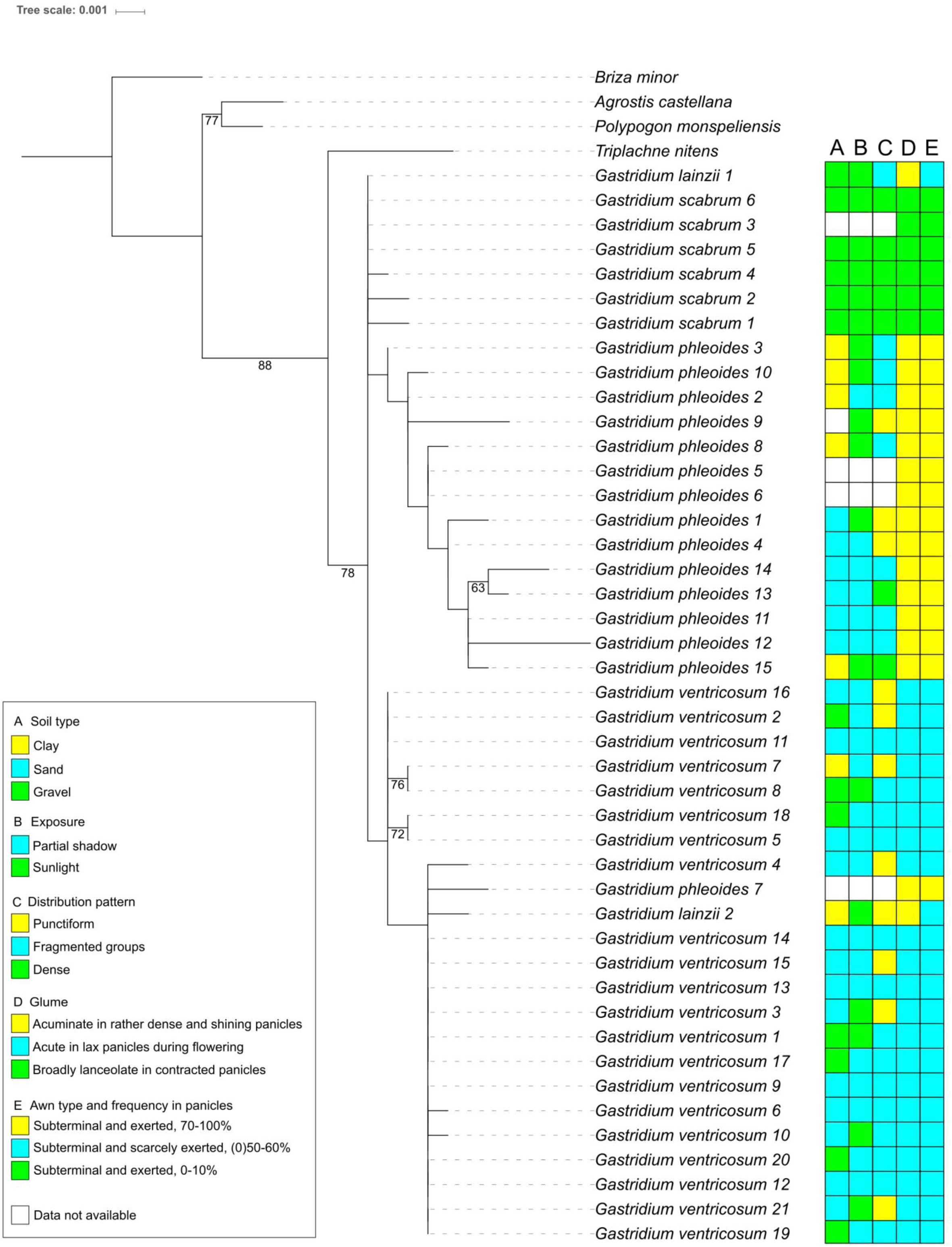
Maximum likelihood phylogram of the trnH-psbA + trnL-F concatenated cpDNA regions in the Agrostidinae dataset. Branch bootstrap support (>50) is reported. Plotted ecological data (soil type, exposure, distribution patterns) were retrieved from field observations and herbarium vouchers; morphological descriptors according to Scoppola & Cancellieri (2019)

The phylogenetic signal of the ITS sequences (Fig. 4) confirmed the poor resolution obtained with the plastid phylogeny. *Agrostis castellana* and *Polypogon monspeliensis* were graded to a highly supported (BS = 100), but unresolved clade that arranged *Gastridium* + *Triplachne*. As in the plastid tree, only a few, minor subclades were weakly or moderately supported, but the groupings had no correspondence with morpho-ecological data or geography, with the exception of the three *G. scabrum* specimens (BS = 67) and two close-by *G. ventricosum* samples (Gv13 and Gv15, BS = 86).

**Fig. 4.**
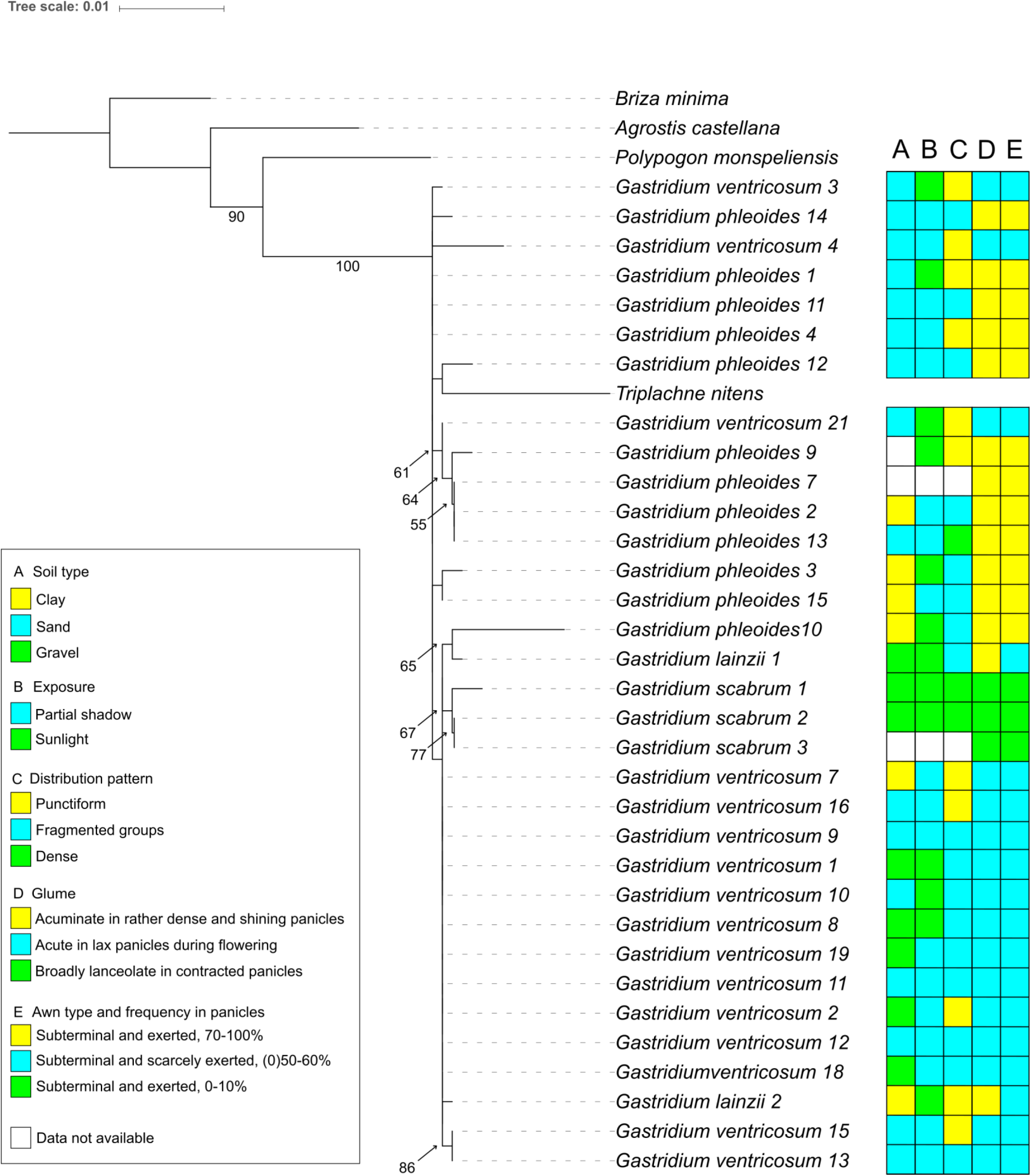
Maximum likelihood phylogram of the nrITS in the Agrostidinae dataset. Branch bootstrap support (>50) is reported. Plotted ecological and morphological data as in Fig. 3

The planar phylogenetic network produced with the Neighbor-Net algorithm provided a clearer representation of the complex phylogenetic relationships produced by the ITS region (Fig. 5). All the outgroup genera (including *Triplachne*) were placed at the tips of long-terminal branches, indicative of high genetic distance with the *Gastridium* dataset (BS = 100). Two major, little supported clusters could be identified, each separately collecting most members of *G. ventricosum* and *G. phleoides.* Samples of both *G. scabrum* and *G. lainzii* split from the *G. ventricosum* sequence core with higher support (57-74). Distinctive placements were recorded for sample Gv21 (placed within the *G. phleoides* cluster), and samples Gv3, Gp15, and Gp3 included in a box-like structure, indicating complex, unresolved relationships. Interestingly, the two *G. lainzii* samples did not display a common or derived phylogenetic origin. Unusual long-terminal branches were scored for samples Gv4 and Gp10.

**Fig. 5.**
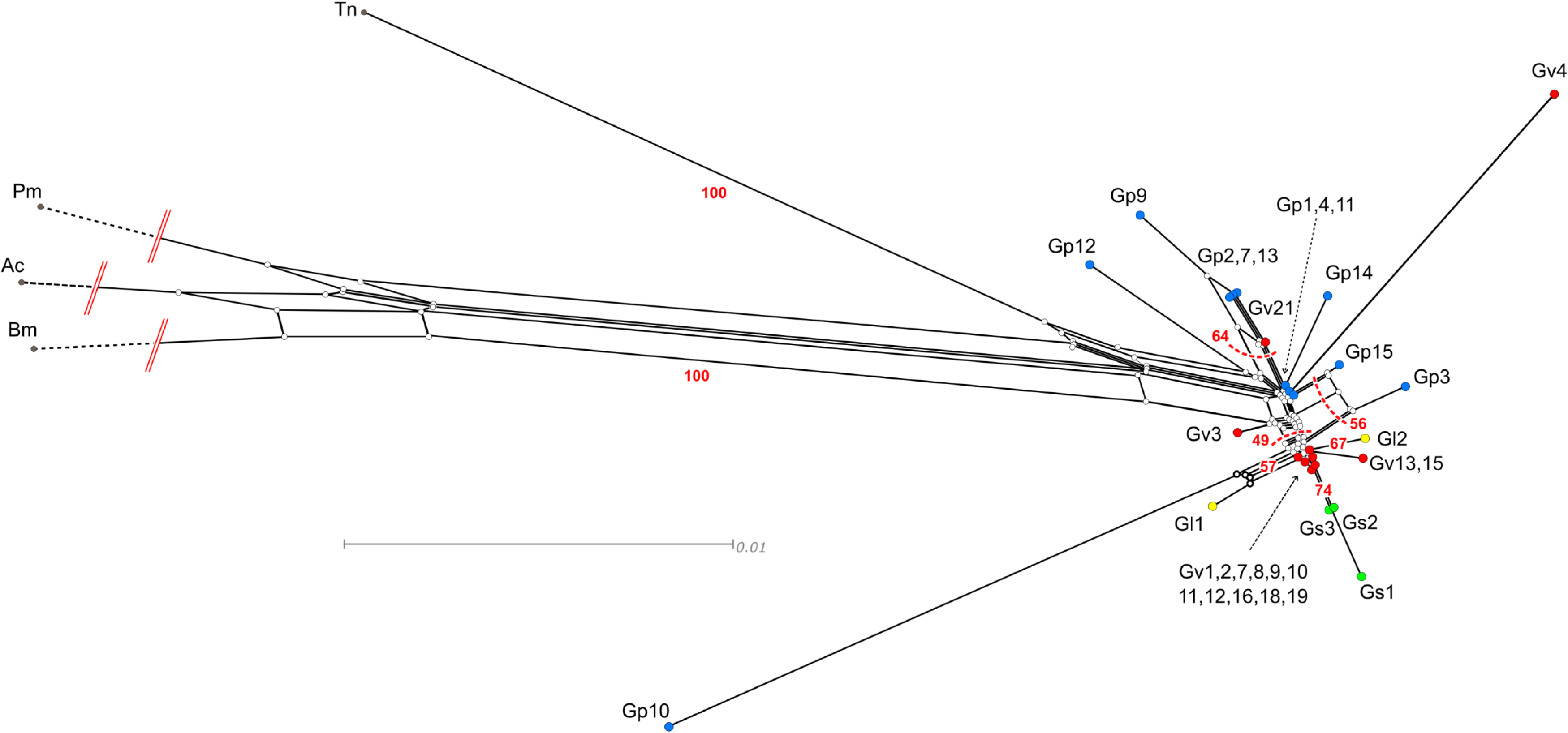
Neighbor-Net splits graph of the nrITS in the Agrostidinae dataset

Integration of our dataset with the trnH-psbA and ITS sequences of members of Agrostidinae currently available in GenBank produced final multiple alignments of 626 and 610 bp, including 189 and 251 sequences, respectively. The resulting RAxML profiles are shown in Figs. 6, 7 (classical rectangular, sequence-labelled ML trees reported as Supplemental online Figure S6a, b). The plastid tree was less resolved than the ITS tree. However, both topologies displayed close relationships and multiple instances of non-monophyly in the analysed genera, together with a general lack of resolution at the species level, largely in agreement with previous phylogenetic reconstructions based on nuclear ITS and more plastid regions (Quintanar et al., 2007; Saarela et al., 2017). In the plastid tree, a sister relationship of *Gastridium* + *Triplachne* (this latter represented by our sample and three sequences retrieved from GenBank) is poorly supported. In the ITS tree, the two genera (including two GenBank sequences of *T. nitens*) formed a highly supported clade but were non-monophyletic. Resolution within the *Gastridium* dataset is lacking in both reconstructions, although sample groupings appear largely congruent with taxa delimitations, especially in the trH-psbA tree.

**Fig. 6.**
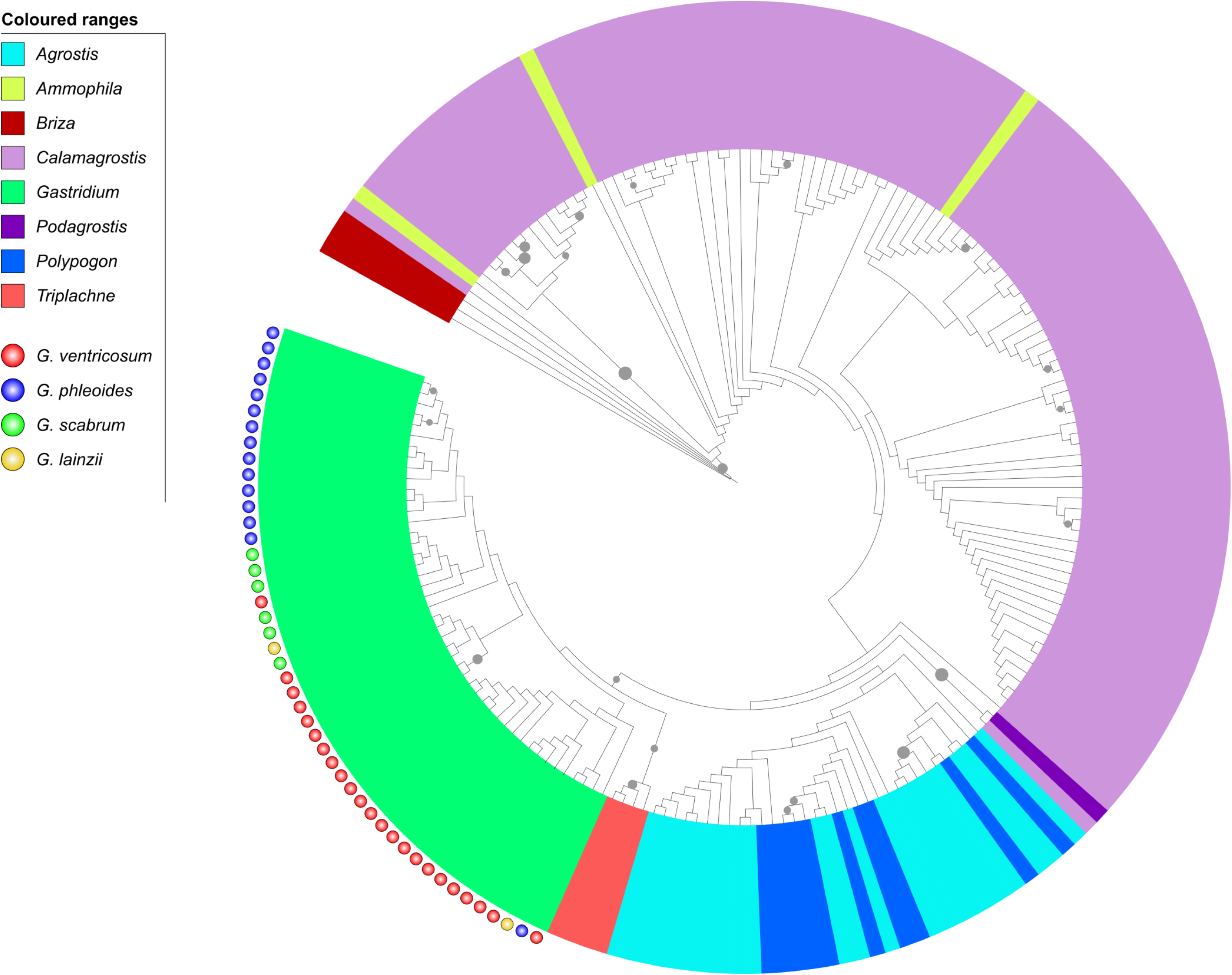
Maximum likelihood trnH-psbA phylogram of the investigated dataset integrated with GenBank sequences of the subtribe Agrostidinae. Grey circles size is proportional to bootstrap support (>50). Coloured ranges indicate different genera. Coloured circles indicate *Gastridium* taxa

**Fig. 7.**
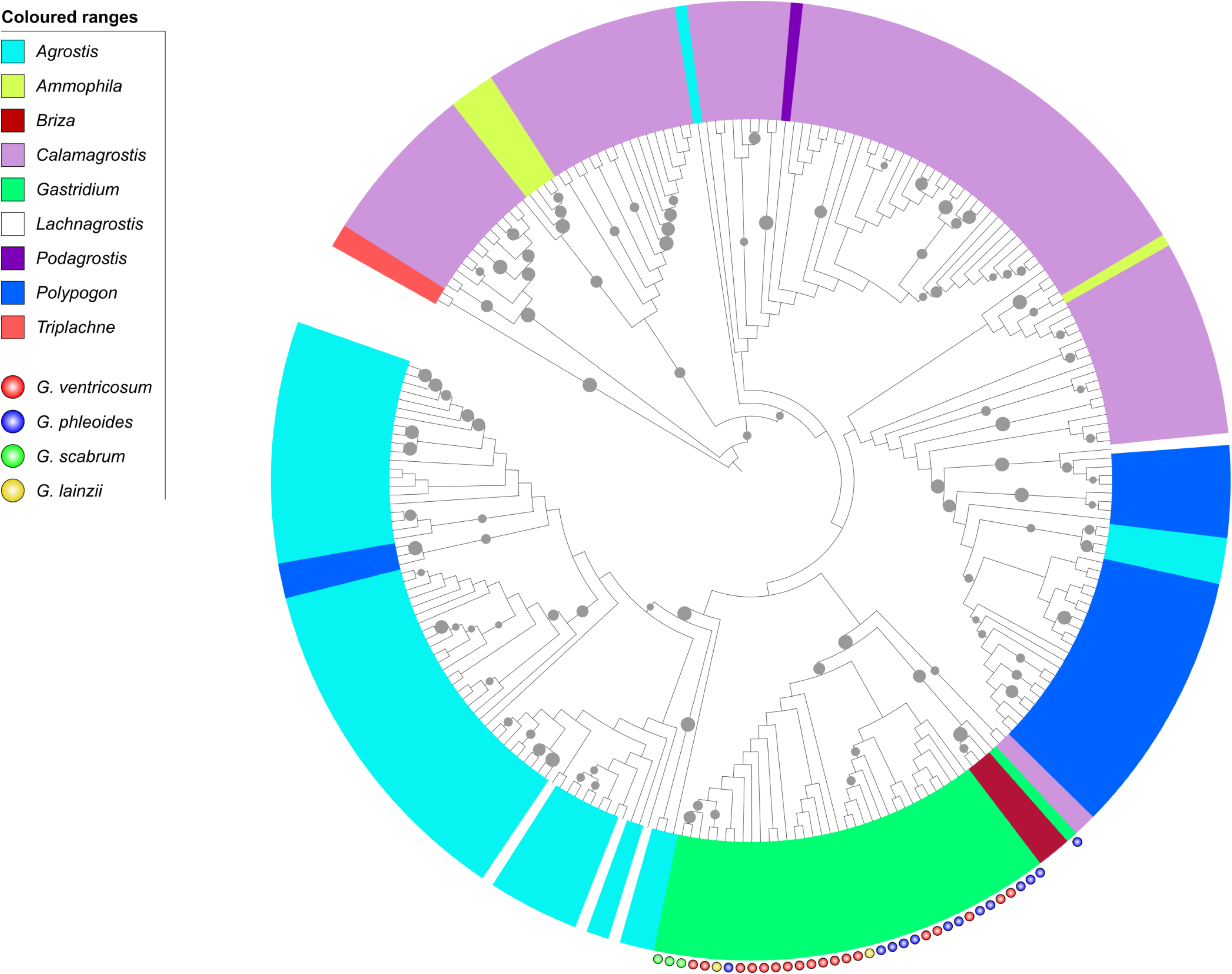
Maximum likelihood nrITS phylogram of the investigated dataset integrated with GenBank sequences of the subtribe Agrostidinae. Grey circles size is proportional to bootstrap support (>50). Coloured ranges indicate different genera. Coloured circles indicate *Gastridium* taxa

Only two trnH-psbA sequences of *G. ventricosum* occurred on GenBank (Acc. Nos. KY432778 and NC036686), but their vouchers were not available for a morphological re-examination. Vouchers corresponding to ITS GenBank sequences DQ336817 (*G. ventricosum*) and KU883502 (*G. phleoides*) revealed the occurrence of the diagnostic traits used in this work in agreement with their assigned names, whereas the sample with sequence KU883503 (*G. ventricosum*) was revised as *G. phleoides*.

## Discussion

### Molecular diversity patterns in *Gastridium*

Our results confirmed the high efficacy of the trnH-psbA to recover diversity patterns at the intrageneric and intraspecific level, especially in complex plant groups where the diagnostic traits are not readily usable (Kress, 2017; Shaw et al., 2005). Despite the limited variation encountered, we found a high occurrence of PICs (Table 1), and the produced haplotypes were not shared among different taxa (File S3). This marker is also known to harbour inversions or deletions that can confound sequence alignments (Whitlock et al., 2010), but we found no evidence of such events in the investigated dataset. In contrast, trnL-F showed less variability, fewer haplotypes with lower diversity, and sequence types shared among different taxa (*G. ventricosum/G. phleoides*, *G. scabrum/G. lainzii*). Nevertheless, the two markers worked synergistically in expanding the diversity levels observed in our dataset. For instance, the 21-bp deletion identified with trnL-F was useful to separate taxa previously treated in synonymy or at subspecific ranks (i.e. *G. scabrum* and *G. lainzii* from *G. ventricosum* and *G. phleoides*), and trnH-psbA further increased their genetic distance. Plastid intergenic spacers are prone to develop high numbers of insertion/deletion mutations (indels) and other microstructural genomic changes with potential phylogenetic utility (Orton et al., 2019). At this regard, no evidence of synapomorphic changes at the inter-/intrageneric level could be identified (cf. Orton et al., 2019), likely due to limited sampling. Likewise, the apomorphic or homoplasious state of the 21-bp indel separating *G. scabrum* and *G. lainzii* from the “major” species cannot be inferred with the available data. Further studies on larger samplings may verify the possible use of this indel as a taxonomic marker for uncertain identifications.

With all markers, *G. ventricosum* and *G. scabrum* were characterized by unambiguous sequence homogeneity (File S4). *Gastridium phleoides* was the most diverse and divergent, and appeared more closely related to *G. ventricosum* with the ITS marker. The only two investigated individuals of *G. lainzii* also were highly diverse and appeared more closely related to *G. ventricosum* and *G. scabrum* at both genomes. However, its sampling was too limited and may have affected the results. Overall, the plastid regions of *G. ventricosum, G. scabrum*, and *G. phleoides* were more divergent than their nuclear ITS. Interestingly, the parallel estimates of the mean intra- and intergroup divergences of the trnH-psbA in these three taxa equalled or exceeded the values recorded in many other species of Agrostidinae. All gathered plastid diversity patterns therefore indicate close relationships among the investigated taxa, a clear distinction of *G. scabrum*, and overall diversity patterns in line with the values recorded within the major lineage they belong. Nevertheless, additional markers and larger samplings would be needed to better assess the extent of plastome variation in *G. lainzii* and enhance our understanding of the high variation displayed by *G. phleoides*.

The ITS marker exhibited higher diversity scores, but the large extent of single mutations found within each sample (mirrored by the low number of PICs encountered) likely indicates that the differences are not yet fixed, and the homogenization process of the intra-group nrDNA arrays is still incomplete (cf. Saarela et al., 2017). In addition, several instances of unreadable electropherograms, likely attributable to intra-individual variation, detection of spurious PCR products and cross-contamination with fungi sequences, resulted in fewer sequences. These drawbacks are well-known (Álvarez & Wendel, 2003; Nieto-Feliner & Rossellò, 2007) and call for special methodological attention and consistent evaluation of the downstream results, making necessary the integration of additional bioinformatics analyses and markers.

### Phylogenetic inferences

The structure of the plastid network (Fig. 2) reflects the low-to-moderate levels of intra- and intergroup differentiation of the *Gastridium* dataset. Nevertheless, although separated by few mutations, haplotypes of different taxa were not intermixed, and specimens sampled more than 1000 km away and assigned to the same taxon were included in the same genetic lineage (e.g., both *G. ventricosum* and *G. phleoides* from Spain and Italy, *G. scabrum* from Italy and Turkey). Likewise, specimens sampled in close proximity and assigned to different taxa clustered separately, in taxonomically uniform lineages (e.g., samples Gv10, Gp1, Gs5, belonging to *G. ventricosum*, *G. phleoides* and *G. scabrum* from the surroundings of Rome; samples Gl1, Gp3, belonging to *G. lainzii* and *G. phleoides* from the surroundings of Cordoba, just to name a few; see Figs. 1, 2). Indeed, geographic patterns of variation can be a relevant information to infer species delimitations and con-specificity (de Queiroz & Good 1997). In this view, our findings support the identification of closely related, separately evolving plastid lineages within *Gastridium*, largely corresponding to the morphologically differentiated taxa and not to geographic patterns of variation. Among the other Agrostidinae samples (*Agrostis castellana*, *Polypogon monspeliensis*, and *Triplachne nitens*), a closer affinity of this latter to the genus *Gastridium* is confirmed, whereas the haplotypes of *Alopecurus aequalis*, *Cynosurus echinatus, Gaudinia fragilis*, and *Briza minor* (all belonging to different subtribes of Pooideae) were too divergent to be adequately evaluated with the network analysis.

In the poorly resolved RAxML plastid tree (Fig. 3), *Gastridium* showed the same sequence partition recorded in the haplotype network, largely corresponding to morphological delimitations, but the identified clusters were unsupported. The *Gastridium* dataset was even less resolved in the ITS tree (Fig. 4). Unresolved tree topologies are well documented across species and genera in Pooideae and grasses in general, and commonly explained with low mutation rates of the used markers, reticulation, polyploidization, hybridization, and lineage sorting (Catalán et al., 2004; Mason-Gamer & Kellogg, 1996; Quintanar et al., 2007; Saarela et al., 2017). Clearly, in our study, the number of PICs detected at both genomes was insufficient to robustly separate the four taxa, and the high number of unfixed mutations in the ITS was scantily deciphered by the RAxML approach. However, it is possible that increasing the number of markers at both genome sources (e.g., Minaya et al., 2015; cf. Saarela et al., 2010, 2017) may enhance the taxonomic resolution here outlined. At the same time, other evolutionary mechanisms may have played an important role in structuring the molecular diversity of *Gastridium*, similarly to other Pooideae lineages (Barberá et al., 2019; Peerson & Rydin, 2016; Soreng et al., 2015b). Past or recent hybridization, for instance, could have been facilitated by the frequent co-occurrence of the investigated taxa (see also the large overlap of ecological descriptors reported in Figs. 3 and 4), and indeed, the diversity patterns observed in *G. lainzii* seem compatible with a hybrid origin (Supplemental online File S4, Fig. 5). In addition, molecular boundaries at both genomes may be masked by polyploidization (Soreng et al., 2017). The ploidy levels of at least two investigated taxa are different (*G. ventricosum* and *G. phleoides* are, respectively, diploid and tetraploid; Kellogg, 2015), whereas *G. scabrum* and *G. lainzii* are still poorly documented, but our preliminary investigations point to a possible tetraploid state for both (Supplemental online material S2).

In general, the low resolution detected in *Gastridium* is not deviating from the data collected in other Agrostidinae genera, where considerable intermixing of species, and little supported species clades, were documented at both the nuclear and the plastid DNA level (c.f. Saarela et al., 2017). This is also confirmed by our expanded trees (Figs. 6, 7), in perfect agreement with large-scale phylogenetic reconstructions obtained with multiple markers (Saarela et al., 2017), or complete plastid genomes (Saarela et al., 2018). Both our reconstructions detected intermixed subsets of species of *Ammophila*, *Agrostis*, *Calamagrostis*, *Lachnagrostis*, and *Polypogon*. The non-monophyly of *Lachnagrostis* and *Ammophila* is well-known (Brown, 2013), and the latter is now included in *Calamagrostis* (Soreng et al., 2017). The close relationship between *Agrostis* and *Polypogon*, their non-monophyly and the species variously intermixed both at the nuclear and the plastid level, are in agreement with Saarela et al. (2017), as well as the low supported sister relationship of these two genera with the *Gastridium* + *Triplachne* clade in the plastid tree.

The clear differentiation of *Gastridium* and *Triplachne* from all other genera and their close relationships recovered at all markers and with all reconstructions reflect the morpho-ecological affinities of these two Mediterranean entities (Clayton & Renvoize, 1986; Pyke, 2008; Scoppola & Cancellieri, 2019). Romero Zarco (2015) even included *Triplachne* within *Gastridium* [*G. nitens* (Guss.) Coss. & Durieu], and recently Romero García (2019) kept them separate, as they differ in some substantial morphological characters (awn position, lemma apex with or without setae, saccate glumes, etc.) and have different habitat preferences. Our data highlight a weakly-supported sister relationship between the two genera at the plastid level, whereas in the ITS tree, only a GenBank sequence of *G. phleoides* (origin: Saudi Arabia) challenged their monophyly. At this regard, we emphasize the higher efficacy of two network approaches (Figs. 2, 5) compared to the classical phylogenetic trees (Figs. 3, 4), in disentangling the complex relationships between these two genera, which are indeed very close but unambiguously divergent.

The ITS Neighbor-Net analysis further contributed to explain the critical position of some variants. For instance, the unusual long-terminal branches scored for samples Gv4 and Gp10 indicate strongly deviating variants, for which it is possible to infer a degenerate (pseudogene) identity (Nieto Feliner & Rossellò 2007). The complex, unresolved relationships linking samples Gv3, Gp15, and Gp3 may indicate reticulation between *G. ventricosum* and *G. phleoides*, while Gv21, Gl1 and Gl2 may subtend hybridization. Cloning or next-generation sequencing approaches would be necessary to understand the nature and origin of the ITS loci in these samples.

Finally, increased sampling can increase phylogenetic information (Zwickl & Hillis, 2002), and conspecific sequences deposited in public databases may provide new insights into intraspecific diversity. In this light, the confirmed identity of specimen with ITS sequence KU883502 (*G. phleoides*) would provide further evidence of the wide genetic diversity existing in this species, and on the complex relationships shared with *Triplachne nitens*, although the homologous nature of the published ITS sequence should first be assessed, followed by further phylogenetic studies and denser regional samplings. At the same time, we remark the importance of an update of the genus current assessment in taxonomic authorities, to circumvent future uncertainties on the diversity of its species.

### Taxonomic implications

Separating closely related species and determining species boundaries is a difficult task, still debated among taxonomists (Pante et al., 2015). Indeed, species can be considered separately evolving metapopulation lineages, where the primary properties usually utilized to define a species (e.g., reproductive isolation, adaptation to specific ecological niches, phenotypic and genetic cohesion, among the others; de Queiroz, 2007) can arise (and become evident, i.e. diagnosable) with different paces during the speciation process (de Queiroz, 1998). Likewise, the states of the characters used for species circumscription pass through polyphyletic, paraphyletic, and monophyletic stages, with a timing depending on ecological, demographic and genetic factors (Avise & Ball, 1990; Funk & Omland, 2003; Hörandl 2006). Recently-derived species are therefore highly prone to show non-monophyletic gene trees, since monophyly will become more detectable as the time since lineage splitting increases (Knowles & Carstens, 2007). At the same time, it is also possible that morphological and genetic differences gradually blur among closely related species, in a sort of continuum (like originally argued by Darwin; Mallet, 2007). In this context, any evidence of lineage differentiation can be of relevance to infer the existence of separate species (de Queiroz, 2007).

In this work, we recovered plastid genealogies consistent with a subdivision of genus *Gastridium* into at least three distinct (Fig. 2), although closely related (Fig. 3), entities (*G. ventricosum, G. phleoides*, and *G. scabrum*). The nuclear ITS showed no (or weak, e.g., the *G. scabrum* minor clade) taxa resolution (Fig. 4) and was indicative of complex relationships, likely originating from reticulation and polyploidization phenomena (Fig. 5; Supplemental online File S2). Both phenomena are very common in the grass family (Soreng et al., 2017); accordingly, plastome analyses can provide increased phylogenetic resolution within subfamilies compared with classical ITS studies (cf. Saarela et al., 2017, 2018).

In this work, *G ventricosum, G. phleoides*, and *G. scabrum* showed evidence of inner genetic cohesion at their plastomes, despite geographic distance among individuals, and inter-taxa mean divergence generally higher than the average estimates within each taxon, despite geographic proximity. In turn, their estimated inter-taxa mean divergences were similar to the interspecific estimates calculated in other genera of the same subtribe. Finally, our plastid phylogenetic reconstructions showed sequence lineages (and evolutionary patterns) largely inclusive of samples sharing diagnosable morphological traits, whereas the ecological descriptors we considered still appear partially overlapping. No substantial difference in the obtained diversity and phylogenetic data seems to justify the different specific ranks assigned to the two acknowledged “major” species (*G. ventricosum*, *G. phleoides*) and *G. scabrum*. Although further investigations are obviously needed, multiple lines of evidence corroborate the hypothesis of progressive lineage separation, likely implying the existence of consolidating, independent species (de Queiroz, 2007). We therefore advance a re-evaluation of the taxonomic treatment of *G scabrum*. This taxon is still treated as a synonym of *G. ventricosum* by the principal taxonomic authorities (Tropicos, GrassBase, IPNI, etc.), although it was validly published in 1818 and shows diagnostic characteristics of the panicle and glumes. In agreement with most authors of Mediterranean regional floras (cf. Doğan, 1985; Feinbrun-Dothan, 1986; Pignatti, 2017; Tison & De Foucault, 2014), our data support its recognition as an independent entity. In addition, preliminary DNA content estimations (Supplemental online material S2) recorded fluorescence intensity scores two times different for *G. ventricosum* and *G. scabrum*, suggesting different ploidy levels in the two entities. It may be worth noting that the Chromosome Counts Database (Rice et al., 2015) treats the two species in synonymy (sub *G. ventricosum*) but enlists mixed counts of 2n = 14 and 2n = 28. De Leonardis et al. (1983) documented a chromosome number = 14 in *G. scabrum*, but the voucher of the investigated specimen is not available for a re-examination.

It is clearly impossible to infer a precise species differentiation pattern with the available data. The low levels of molecular diversity detected, as well as the loose morpho-ecological boundaries within the genus, likely suggest recent diversification. We may speculate that the plastid phylogenetic reconstructions and the known ploidy level (2n = 14) point at *G. ventricosum* as the primitive species. Its most likely ancestral habitats could have been the ephemeral and fragmented grass patches and fringes in meso-Mediterranean open forests and garrigues, characterized by annual, poorly competitive species, where it still occurs (cf. Gallagher et al., 2019). Its limited genetic diversity likely reflects poor adaptability or progressive erosion. Subsequent climate changes and/or human-induced open habitats (pathways, shrubby pastures, hedges) may have subsequently triggered species differentiation, promoting adaptation to drier conditions and disturbance (Clayton & Renvoize, 1986). Among the derived species, all adapted to dry Mediterranean habitats, the wider ecological niche and distribution range of *G. phleoides* (2n = 28) likely indicate a stronger ability of this species to cope with disturbance, and higher dispersal and reproductive efficiency (Lovat, 1981). This would be in agreement with the high genetic diversity detected with both genomes‟ markers, and the most substantial variation scored in the characters of its florets (Scoppola & Cancellieri, 2019). In contrast, the relative homogeneity found in *G. scabrum* matches its limited morpho-ecological variability and may be explained with a recent derivation (by polyploidization) of *G. ventricosum*. Finally, little can be said about *G. lainzii*, only recently proposed as a separate species and previously described as a subspecies of *G. phleoides* (Romero García, 1996). Interestingly, despite the limited sampling and restricted distribution range, this taxon exhibited high genetic diversity, and appeared more closely related to *G. ventricosum*. It may therefore represent a recently diverged lineage of this species, more adapted to drier habitats at the south-western limits of its range. A hybrid origin of this taxon should not be discarded, however, and it clearly deserves further investigations.

## Conclusion

In this work, we took advantage of a recent and upgraded morphometric study of *Gastridium* to expand the individual number of clearly differentiated taxa investigated in a phylogenetic context. Diversity patterns at the nuclear and plastid DNA level, and the different phylogenetic signal of the two analysed genomes were evaluated to reconcile molecular evidence, morphology, and taxonomic congruence. Our results confirm that molecular data can be a useful complement to taxonomic definitions in complex or poorly known taxonomic groups. Although increased sampling efforts and investigations with additional markers would be obviously needed to better understand the complex relationships among the investigated taxa, our results demonstrate that the morphological diversity in *Gastridium* has consistent molecular grounds in the plastid and nuclear genomes, and likely subtend the existence of more species than currently acknowledged. Together with *G. ventricosum, G. phleoides* and the recently acknowledged *G. lainzii*, we argue that *G. scabrum* also deserves species status. These species may constitute a set of recent origin where the morphological and genetic differences are evident, although only partly fixed and still limited. In this view, the role played by polyploidization and hybridization in the genus diversification needs to be firstly assessed. Unambiguous information on the systematics and phylogeny of genera is a fundamental prerequisite to biodiversity evaluation, ecosystem conservation, and the improvement of our knowledge of the tree of life (Mace, 2004). The presented data provide hints for a reassessment of the taxonomy of *Gastridium*, to improve research and protection of ephemeral habitats of the Mediterranean in a context of global change.

## Acknowledgements

We gratefully acknowledge the Directors and Curators of MA, SBT, SNM, UPS, and quoted herbaria for providing plant material and scanned images of herbarium specimens. Thanks are due to the U.S. National Plant Germplasm System for providing *G. scabrum* caryopses, J.A. Devesa Alcaraz (Cordoba) for the sample of *T. nitens*, P. Catalán (Zaragoza), O. Ryding (Copenhagen), E. Lattanzi and F. Lucchese (Roma) for useful information, R. Tenchini (Viterbo) for technical assistance. This research was supported by MIUR (Ministry for Education, University and Research), Law 232/2016, “Department of excellence” and the “Cordero Lanza di Montezemolo” Donation.

## Supplement

**Supplementary File S1.** This file contains all relevant data concerning the investigated dataset.

**Supplementary File S2.** Flow cytometry analysis

**Supplementary File S3.** List of the plastid haplotypes and nuclear ITS variants.

**Supplementary File S4**. This file contains heatmaps.

**Supplementary File S5a.** TrnL-F haplotype network of the investigated dataset.

**Supplementary File S5b.** TrnH-psbA haplotype network of the investigated dataset.

**Supplementary File S6a.** Maximum likelihood phylogram inferred from the plastid trnH-psbA with 189 sequences of the subtribe Agrostidinae retrieved from GenBank.

**Supplementary File S6b.** Maximum likelihood phylogram inferred from nrITS with 251 sequences of the subtribe Agrostidinae retrieved from GenBank.

