## Supplementary File S2 for "Genetic diversity and taxonomic issues in *Gastridium* P.Beauv (Poaceae) inferred from plastid and nuclear DNA sequence analysis"

**Supplementary File S2 – Flow cytometry analysis**

Determination of the DNA content of a species provides important information for research on molecular and cell genetics and the evolution of plant genomes. Chromosome counting in single plant cells is the traditional method for determining ploidy level; however, this method is time-demanding and laborious, especially when little information is known on the target species (Doležel et al. 2007). Flow cytometry is a fast and accurate, widely used method for the estimation of plant nuclear genome sizes, and differences in DNA content between species are scored by the relative fluorescence intensity of isolated nuclei stained with a DNA-specific fluorochrome. DNA content estimates can be then be used for comparison among conspecifics for routine ploidy estimation, or to standards, in case determination of precise DNA amounts is pursued (Doležel et al. 2007).

### Materials and Methods

We compared the relative fluorescence intensity emitted by 2-3 plants of *G. ventricosum* (2n = 14), *G. phleoides* (2n = 28) (Kellogg 2015), *G. scabrum* and *G. lainzii* (ploidy level unknown), both collected in the field and grown in pots (origin of the samples provided in Table 1). Approximately, 20 mg of fresh and healthy leaves were chopped in Petri dishes (in ice) containing 1ml LB01 lysis buffer (5mM Tris, 2mM Na2EDTA, 0.5mM spermine 4 HCl, 80mM KCl, 20mM NaCl, 0.1% TritonX-100, pH 7.5; Doležel et al. 1989). The homogenate was incubated for 10 minutes with RNAse on ice, filtered with 30µm filters, then 100 µl propidium iodide staining solution (100mg/l) were added. Samples were analysed on a NovoCyte Flow Cytometer (Bioscience, inc.) after 30 min incubation on ice, and the flow cytometric data were collected with the NovoExpress 1.3.0 software.

**Table 1**. Dataset and main results of the Flow Cytometry analysis

| Taxon | Origin | Collection | Voucher | Mean x | CVx |
|---|---|---|---|---|---|
| *G. ventricosum* | Italy (Tuscania) | Wild | UTV-35254 | 181,113 | 4.78 % |
| *G. ventricosum* | Italy (Barbarano Romano) | Wild | UTV-22518 | 137,064 | 6.05 % |
| *G. ventricosum* | Italy (Bomarzo) | Wild | UTV-35250 | 159,523 | 4.81 % |
| *G. phleoides* | Italy (Ischia di Castro) | Wild | UTV-33935 | 350,612 | 3.49 % |
| *G. phleoides* | Italy (Ischia di Castro) | Wild | UTV-35480 | 377,294 | 3.81 % |
| *G. phleoides* | Italy (Bracciano) | Wild | UTV-35577 | X | X |
| *G. scabrum* | Turkey (Izmir) | Pot | UTV-37841 | 418,026 | 2.23 % |
| *G. scabrum* | Italy (Tolfa) | Wild | UTV-13684 | 416,504 | 4.05 % |
| *G. scabrum* | Italy (S. Marinella) | Wild | UTV-35261 | X | X |
| *G. lainzii* | Spain (Cadiz) | Pot | UTV-35257 | 350,757 | 3.04 % |
| *G. lainzii* | Spain (Cordoba) | Pot | UTV-35581 | 386,050 | 4.69 % |

Mean x: relative fluorescence of the stained nuclei; CVx: coefficient of variation of the fluorescent peaks

### Results

Overall, we counted between 200 and 1400 nuclei in each sample, with coefficient of variation (CV) ranging between 2.23 and 6.05%. Best (more consistent) results were obtained with young leaves. In contrast, older leaves either collected from samples in the wild or grown in pots provided less nuclei, largely arrested in G2 phase and more debris; these samples were filtered from the analysis. Although the quality of the analysis was generally limited by some technical problems (low sample numbers, high debris background, low number of G1 isolated nuclei), mostly related with the lack of standard methods for the target taxa and the scarce availability of samples for identifying the most proper plant stage, our measurements indicated (Tab. 1) that the relative nuclear DNA fluorescence intensity of *G. ventricosum* (2n = 14) and *G. phleoides* (2n = 28) are clearly two times different (159,523-181,113 and 350,612-377,294, respectively; only samples with CVx<5% considered), in agreement with their known ploidy level. Illustrative examples of the two species are reported in Fig. 1-2.

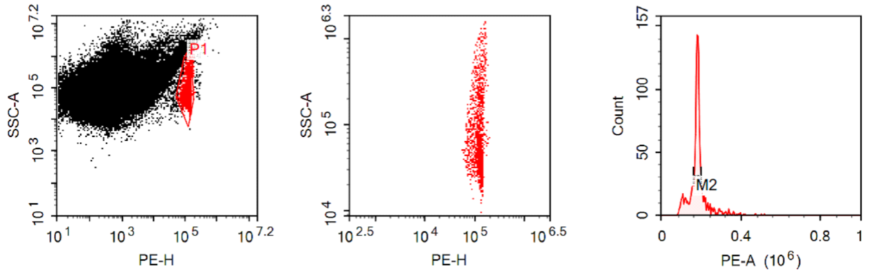


| Gate | Count | % P1 | Mean X | CV X |
|---|---|---|---|---|
| P1 | 1,266 | 100.00 % | 184,052 | 26.21 % |
| M2 | 814 | 64.30 % | 181,113 | 4.78 % |

Sample Statistics of GV1 tris

| Gate | Color | Count | % Parent | X | Y | Mean X | Mean Y | CV X | CV Y |
|---|---|---|---|---|---|---|---|---|---|
| All | 3 | 100,000 | | | | | | | |
| └ P1 | 1 | 1,266 | 1.27 % | PE-H | SSC-A | 118,256 | 157,672 | 19.85 % | 148.35 % |
| └ M2 | 2 | 814 | 64.30 % | PE-A | Count | 181,113 | | 4.78 % | |

Fig. 1 – Results report *of G. ventricosum* from Tuscania

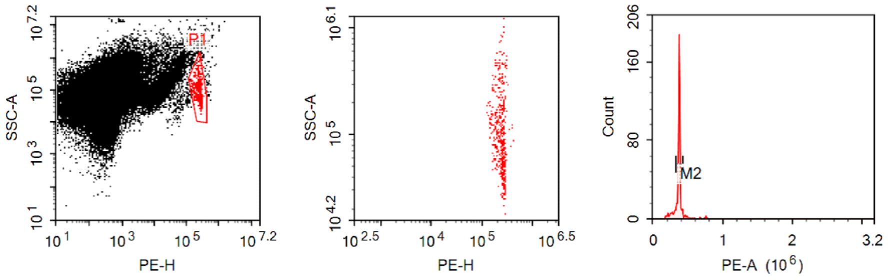


| Gate | Count | % P1 | Mean X |
|---|---|---|---|
| P1 | 480 | 100.00 % | 376,041 |
| M2 | 372 | 77.50 % | 377,294 |

Sample Statistics of GP2 tris

| Gate | Color | Count | % Parent | X | Y | Mean X | Mean Y | CV X | CV Y |
|---|---|---|---|---|---|---|---|---|---|
| All | 3 | 100,000 | | | | | | | |
| └ P1 | 1 | 480 | 0.48 % | PE-H | SSC-A | 237,736 | 143,915 | 17.65 % | 107.79 % |
| └ M2 | 2 | 372 | 77.50 % | PE-A | Count | 377,294 | | 3.81 % | |

Fig. 2 – Results report of *G. phleoides* from Ischia di Castro, sample 2

*Gastridium scabrum* and *G. lainzii* deviated sensibly from the *G. ventricosum* fluorescence intensity records; rather they were more similar to the values scored by *G. phleoides*, which makes possible to assume a tetraploid state for both (Fig. 3).

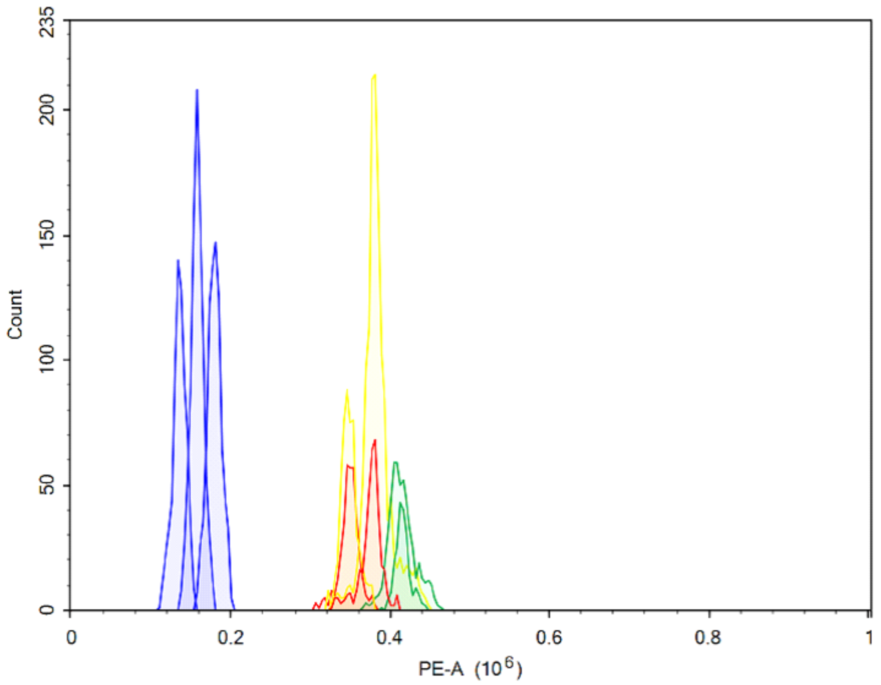


Fig. 3 – Relative fluorescence intensity measured on the *Gastridium* dataset. Blue: *G. ventricosum*; red: *G. phleoides*, green: *G. scabrum*; yellow: *G. lainzii*

Indeed, further studies are required to precisely assess DNA contents and ploidy levels in *Gastridium*. Nevertheless, differentiation of *G. ventricosum* from the three other taxa (and especially from *G. scabrum*, traditionally treated as a synonym or a subspecies of *G. ventricosum*) seemed evident. In future studies, one useful strategy to overcome the encountered drawbacks would be to analyse only plants growing under controlled conditions (3-5 samples per accession is advisable) to identify the most suitable leaf stage and testing different nuclei isolation buffers.
