## Supplementary File S3 for "Genetic diversity and taxonomic issues in *Gastridium* P.Beauv (Poaceae) inferred from plastid and nuclear DNA sequence analysis"

**Supplementary File S3**: List of the plastid haplotypes and nuclear ITS variants detected in the investigated dataset; sequence codes according to Supplementary File S1.

| **TrnH-psbA** | | |
| --- | --- | --- |
| Haplotype | Frequency | Sequences |
| Hap_1 | 13 | Gv1 - Gv3 - Gv4 - Gv6 - Gv9 - Gv12 - Gv13 - Gv14 - Gv15 - Gv17 - Gv19 - Gv20 - Gv21 |
| Hap_2 | 5 | Gv2 - Gv5 - Gv11 - Gv16 - Gv18 |
| Hap_3 | 2 | Gv7 - Gv8 |
| Hap_4 | 1 | Gv10 |
| Hap_5 | 1 | Gp1 |
| Hap_6 | 1 | Gp2 |
| Hap_7 | 1 | Gp3 |
| Hap_8 | 3 | Gp4 - Gp5 - Gp6 |
| Hap_9 | 1 | Gp7 |
| Hap_10 | 1 | Gp8 |
| Hap_11 | 1 | Gp9 |
| Hap_12 | 1 | Gp10 |
| Hap_13 | 2 | Gp11 - Gp12 |
| Hap_14 | 1 | Gp13 |
| Hap_15 | 1 | Gp14 |
| Hap_16 | 1 | Gp15 |
| Hap_17 | 5 | Gs1 - Gs3 - Gs4 - Gs5 - Gs6 |
| Hap_18 | 1 | Gs2 |
| Hap_19 | 1 | Gl1 |
| Hap_20 | 1 | Gl2 |
| Hap_21 | 1 | Tn |
| Hap_22 | 1 | Pm |
| Hap_23 | 1 | Ac |
| Hap_24 | 1 | Bm |
| Hap_25 | 1 | Ae |
| Hap_26 | 1 | Ce |
| Hap_27 | 2 | Gf - Gf2 |

| **TrnL- TrnF** | | |
| --- | --- | --- |
| Haplotype | Frequency | Sequences |
| Hap_1 | 24 | Gv1 - Gv2 - Gv3 - Gv7 - Gv8 - Gv9 - Gv10 - Gv11 - Gv12 - Gv13 - Gv14 - Gv15 - Gv16 - Gv17 Gv19 - Gv20 - Gv21 - Gp2 - Gp3 - Gp5 - Gp6 - Gp8 - Gp9 - Gp10 |
| Hap_2 | 1 | Gv4 |
| Hap_3 | 2 | Gv5 - Gv18 |
| Hap_4 | 1 | Gv6 |
| Hap_5 | 6 | Gp1 - Gp4 - Gp11 - Gp13 - Gp14 - Gp15 |
| Hap_6 | 1 | Gp7 |
| Hap_7 | 1 | Gp12 |
| Hap_8 | 1 | Gs1 |
| Hap_9 | 6 | Gs2 - Gs3 - Gs5 - Gs6 - Gl1 - Gl2 |
| Hap_10 | 1 | Gs4 |
| Hap_11 | 1 | Tn |
| Hap_12 | 1 | Pm |
| Hap_13 | 1 | Ac |
| Hap_14 | 1 | Bm |
| Hap_15 | 1 | Ae |
| Hap_16 | 1 | Ce |
| Hap_17 | 2 | Gf - Gf2 |

| **TrnH-psbA + TrnL- TrnF** | | |
| --- | --- | --- |
| Haplotype | Frequency | Sequences |
| Hap_1 | 11 | Gv1 - Gv3 - Gv9 - Gv12 - Gv13 - Gv14 - Gv15 - Gv17 - Gv19 - Gv20 - Gv21 |
| Hap_2 | 3 | Gv2 - Gv11 - Gv16 |
| Hap_3 | 1 | Gv4 |
| Hap_4 | 2 | Gv5 - Gv18 |
| Hap_5 | 2 | Gv7 - Gv8 |
| Hap_6 | 1 | Gv6 |
| Hap_7 | 1 | Gv10 |
| Hap_8 | 1 | Gp1 |
| Hap_9 | 1 | Gp2 |
| Hap_10 | 1 | Gp3 |
| Hap_11 | 1 | Gp4 |
| Hap_12 | 2 | Gp5 - Gp6 |
| Hap_13 | 1 | Gp7 |
| Hap_14 | 1 | Gp8 |
| Hap_15 | 1 | Gp9 |
| Hap_16 | 1 | Gp10 |
| Hap_17 | 1 | Gp11 |
| Hap_18 | 1 | Gp12 |
| Hap_19 | 1 | Gp13 |
| Hap_20 | 1 | Gp14 |
| Hap_21 | 1 | Gp15 |
| Hap_22 | 1 | Gs1 |
| Hap_23 | 1 | Gs2 |
| Hap_24 | 3 | Gs3 - Gs5 - Gs6 |
| Hap_25 | 1 | Gs4 |
| Hap_26 | 1 | Gl1 |
| Hap_27 | 1 | Gl2 |
| Hap_28 | 1 | Tn |
| Hap_29 | 1 | Pm |
| Hap_30 | 1 | Ac |
| Hap_31 | 1 | Bm |
| Hap_32 | 1 | Ae |
| Hap_33 | 1 | Ce |
| Hap_34 | 2 | Gf - Gf2 |

| **ITS** | | |
| --- | --- | --- |
| Haplotype | Frequency | Sequences |
| Hap_1 | 11 | Gv1 - Gv2 - Gv7 - Gv8 - Gv9 - Gv10 - Gv11 - Gv12 - Gv16 - Gv18 - Gv19 |
| Hap_2 | 1 | Gv3 |
| Hap_3 | 1 | Gv4 |
| Hap_4 | 2 | Gv13 - Gv15 |
| Hap_5 | 1 | Gv21 |
| Hap_6 | 3 | Gp1 - Gp4 - Gp11 |
| Hap_7 | 3 | Gp2 - Gp7 - Gp13 |
| Hap_8 | 1 | Gp3 |
| Hap_9 | 1 | Gp9 |
| Hap_10 | 1 | Gp10 |
| Hap_11 | 1 | Gp12 |
| Hap_12 | 1 | Gp14 |
| Hap_13 | 1 | Gp15 |
| Hap_14 | 1 | Gs1 |
| Hap_15 | 2 | Gs2 - Gs3 |
| Hap_16 | 1 | Gl1 |
| Hap_17 | 1 | Gl2 |
| Hap_18 | 1 | Tn |
| Hap_19 | 1 | Pm |
| Hap_20 | 1 | Ac |
| Hap_21 | 1 | Bm |
| Hap_22 | 1 | Ae |
| Hap_23 | 1 | Ce |
| Hap_24 | 1 | Gf |
