## Supplementary File S5b for "Genetic diversity and taxonomic issues in *Gastridium* P.Beauv (Poaceae) inferred from plastid and nuclear DNA sequence analysis"

**Supplementary File S5b.** TrnH-psbA haplotype network of the investigated dataset. Coloured ellipses circumscribe the four investigated *Gastridium* taxa. Taxa colours as in Fig.1. Different line thickness indicates the relative number of mutations. *Ce: Cynosurus echinatus, Ae: Alopecurus aequalis, Gf: Gaudinia fragilis, Bm: Briza minor, Ac: Agrostis castellana, Pm: Polypogon monspeliensis, Tn: Triplachne nitens, Gv: Gastridium ventricosum, Gp: G. phleoides, Gs: G. scabrum, Gl: G. lainzii.*

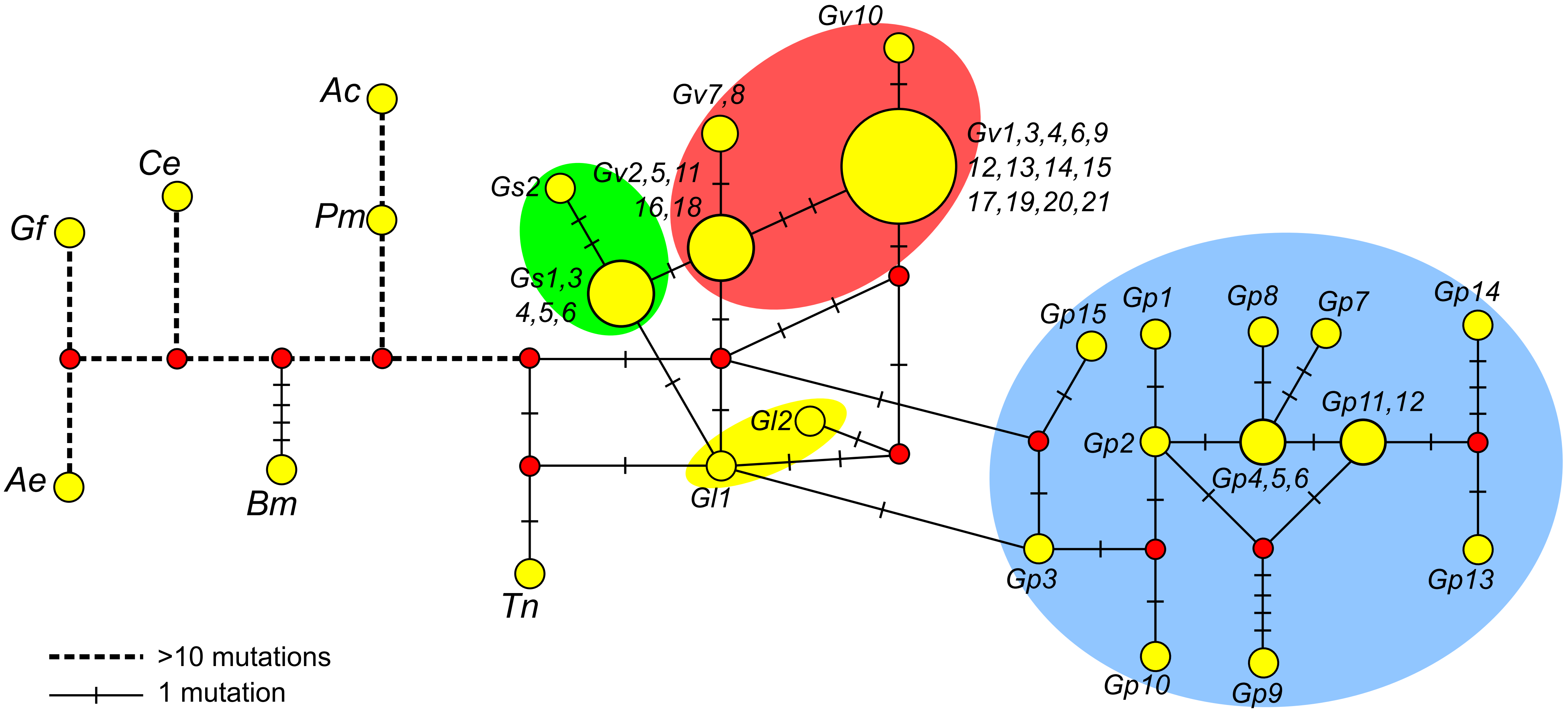
