## Supplementary File S6b for "Genetic diversity and taxonomic issues in *Gastridium* P.Beauv (Poaceae) inferred from plastid and nuclear DNA sequence analysis"

**Supplementary File S6b.** Maximum likelihood phylogram inferred from nrITS of the investigated dataset integrated with 251 sequences of the subtribe Agrostidinae (Soreng et al. 2017) retrieved from GenBank. Bootstrap support (>50) is indicated on the corresponding branches. Colouration refers to different genera.

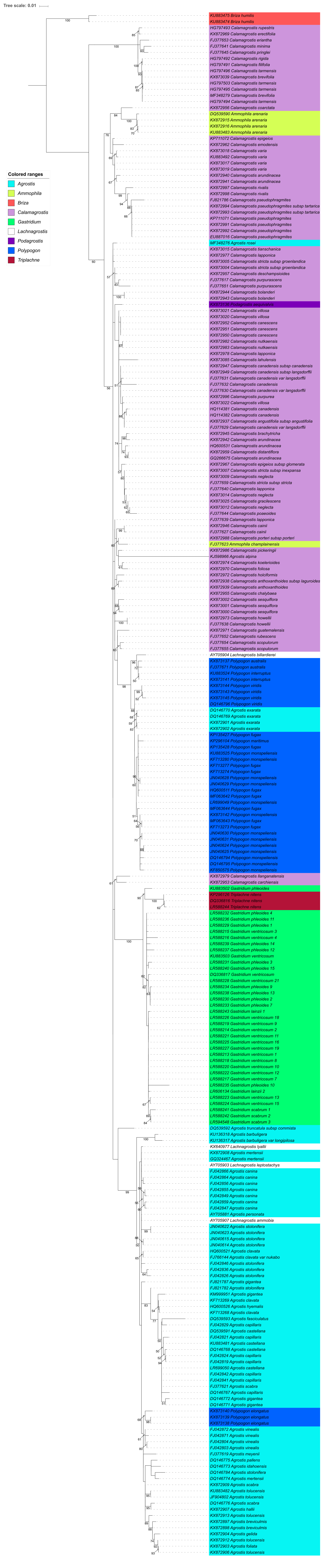
